# An explainable artificial intelligence-based typification of chronic inflammatory responses enhances glioma prognosis

**DOI:** 10.1101/2023.02.28.530381

**Authors:** Debajyoti Chowdhury, Hiu Fung Yip, Zeming Li, Qing Ren, Hao Liu, Xuecheng Tai, Lu Zhang, Aiping Lu

## Abstract

Glioma is one of the most aggressive solid brain tumors with a poor prognosis. A chronic tumor inflammatory microenvironment drives glioma promotion and progression. The neutrophil-to-lymphocyte ratio and other clinicopathological variables usually serve as prognostic glioma markers. However, they are not ubiquitous prognostic markers for glioma as they fail to reveal the intricacy between the glioma-specific tumor inflammatory microenvironment and the systemic inflammatory responses, especially those chronic inflammatory responses, which vary among individuals fabricating diverse prognostic outcomes. Here, we introduced an explainable artificial intelligence model to typify chronic inflammatory responses as prognostic markers for glioma using 694-patients’ data from The Cancer Genome Atlas. We characterized the glioma-specific personalized inflammatory mediators using multi-layered regulators such as transcriptional networks, cellular infiltration markers, and cellular senescence markers, which identified five unique chronic inflammatory responses (p-value<0.0001). We defined its prognostic significance using overall survival analyses. The chronic inflammatory responses were positively correlated with poor overall survival in glioma. The patients with higher chronic inflammatory responses showed significantly shorter overall survival than those with lower chronic inflammatory responses. Interestingly, optimizing those chronic inflammatory responses improved the overall survival of glioma patients. We identified the effector genes within the personalized inflammatory mediators’ networks, indicating them as the targets for optimizing individualized chronic inflammatory response profiles through co-drug intervention.

**Significance:** Explainable artificial intelligence-based typification of chronic inflammatory responses accelerates glioma prognosis and supports co-drug discovery to modulate inflammatory responses alongside cancer therapy, suggested by 694-glioma patients’ data analysis.

## 1. Introduction

Despite the human immune system’s adaptiveness in responding to various potential threats, unnecessary and uncontrolled inflammatory responses cause various chronic illnesses, including cancers(1) (Figure 1A). Nearly 20% of cancer-related casualties are inflammatory responses related(2), which is not extensively evaluated with the existing drug discovery pipeline and causes a massive financial loss(3). Furthermore, these under-characterized chronic inflammatory responses (CIRs) fluctuate among cancer patients affecting prognostic assessments(4,5).

**Figure 1:**
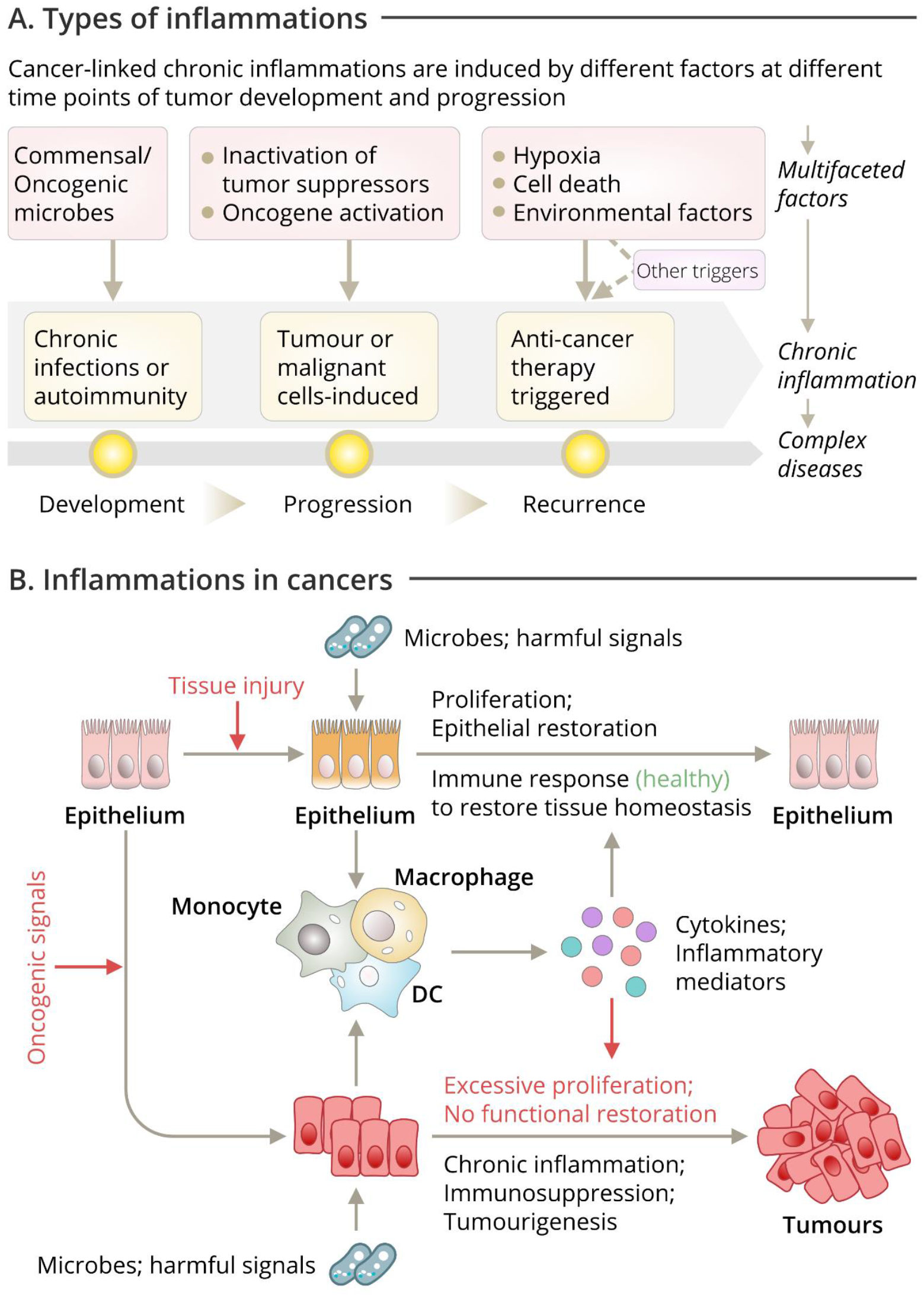
Overview of the types of inflammations in cancers. (A) Cancer-linked chronic inflammations induced by different factors at different time point of the disease progression. (B) Inflammatory response generation mechanism in cancer at cellular level.

Glioma, the most common and aggressive form of solid brain tumors with a poor prognosis, is substantially influenced by uncontrolled CIRs(6,7). The glioma-specific tumor inflammatory microenvironment (TIME) comprises various inflammatory mediators and cellular effectors(7) that drive its development, promotion, and progression upon altering the steady immune systems(1) (Figure 1A, 1B). CIRs usually show two distinct roles in cancers, one, they may exist before any malignant changes occur; two, the oncogenic modifications may impair steady TIME (2,8). In addition, the CIRs used to induce angiogenesis and metastasis by destabilizing the adaptive immune responses, which further constricts therapeutic responses(9) in glioma. Details of those CIRs are not yet well illustrated, leading to various clinical challenges(10). The neutrophil-to-lymphocyte ratio (NLR) combined with pathological variables is commonly used as prognostic markers (PMs) for glioma(11). But they do not adequately expose the intricacy between the glioma TIME and the systemic CIRs. So, they cannot be considered as the ubiquitous PM for glioma, including both glioblastoma (GBM) and lower-grade glioma (LGG). Furthermore, many chemotherapies are constrained due to poor management of the concurrent CIRs varying across individuals(11,12). So, it is sensitive and challenging to opt the accurate chemotherapy plans. The key challenges include an underdetermined cellular origin of gliomas(13), a lack of characterization of CIRs acting differently across glioma patients(9), and a lack of strategic prognosis and personalized therapeutic recommendations(14,15).

Here, we introduced an explainable artificial intelligence (XAI) model that typified the CIRs into five unique rational classes upon analyzing the whole transcriptome of CIRs-related signatures from 694 glioma patients and improved prognostic assessment. The model is flexible to switch between population-based and personalized prognosis. We have also shown a strategic co-drug discovery pipeline to modulate those CIRs. It eventually intends to provoke a reimagined therapeutic co-intervention strategy for glioma where the co-drugs can be recommended along with primary chemotherapy to mitigate the unnecessary concurrent CIRs without interfering.

## 2. Results

We have developed an XAI model that identifies the personalized inflammatory mediators (PIMs) to characterize and rationalize the CIRs serving as efficient PMs for glioma. We typified five unique CIR types at p-value<0.0001 (Figure 2B). The prognostic ability assessed with overall survival (OS) analyses indicated the higher level of CIRs showed significantly lower OS and vice versa for the glioma population, GBM at p-value<0.22 and LGG at p-value<0.0001 (Figure 2C). We showed that the CIR type 2 and type 4 were highly inflamed and had poor OS for GBM, and LGG, respectively. Our results also outperformed the conventional NLR-based glioma prognosis (Figure 2D). As the extent of CIRs was showed positive correlation with poor OS for glioma patients, optimizing the CIRs may improve the OS for glioma patients. Then, we identified the essential driver genes (EDGs) those critically determine the TIME for the GBM and LGG patients (Figure 3A). We also identified the personalized EDGs (pEDGs) within indivudal’s CIR network, which served as potential targets for modulating individual CIRs through co-drug interventions (Figure 4).

**Figure 2:**
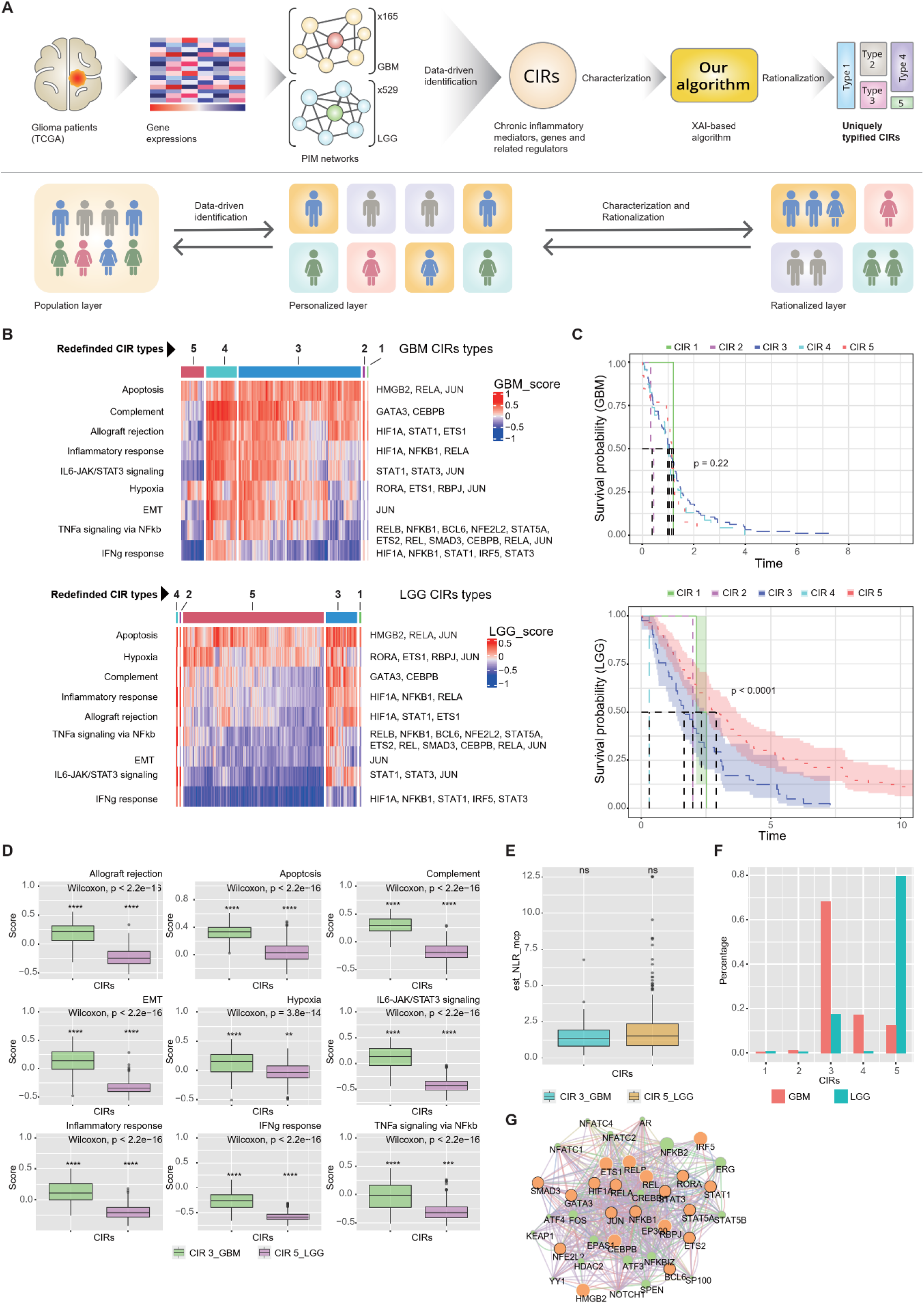
The overview of the proposed XAI model and its typification performance. (A) The XAI model-based analysis pipeline. (B) The GSVA activation score of the top nine enriched hallmark pathways and the corresponding heatmap for GBM (upper panel) and LGG (lower panel). (C) The Kaplan-Meier curves of overall survival with Log-Rank p-value for the typified CIRs type in gliomas, GBM (upper panel) and LGG (lower panel). (D) Box plots of the GSVA activation scores for those nine enriched pathways for GBM CIR type 3 and LGG CIR type 5 with Wilcoxon’s test. (E) Box plot of NLR score for GBM CIR type 3 and LGG CIR type 5 with Wilcoxon’s test. (F) Bar graph showing the percentage distribution of the five CIRs types in GBM and LGG. (G) The transription factors networks identified from the nine enriched hallmark pathways using GeneMANIA.

**Figure 3:**
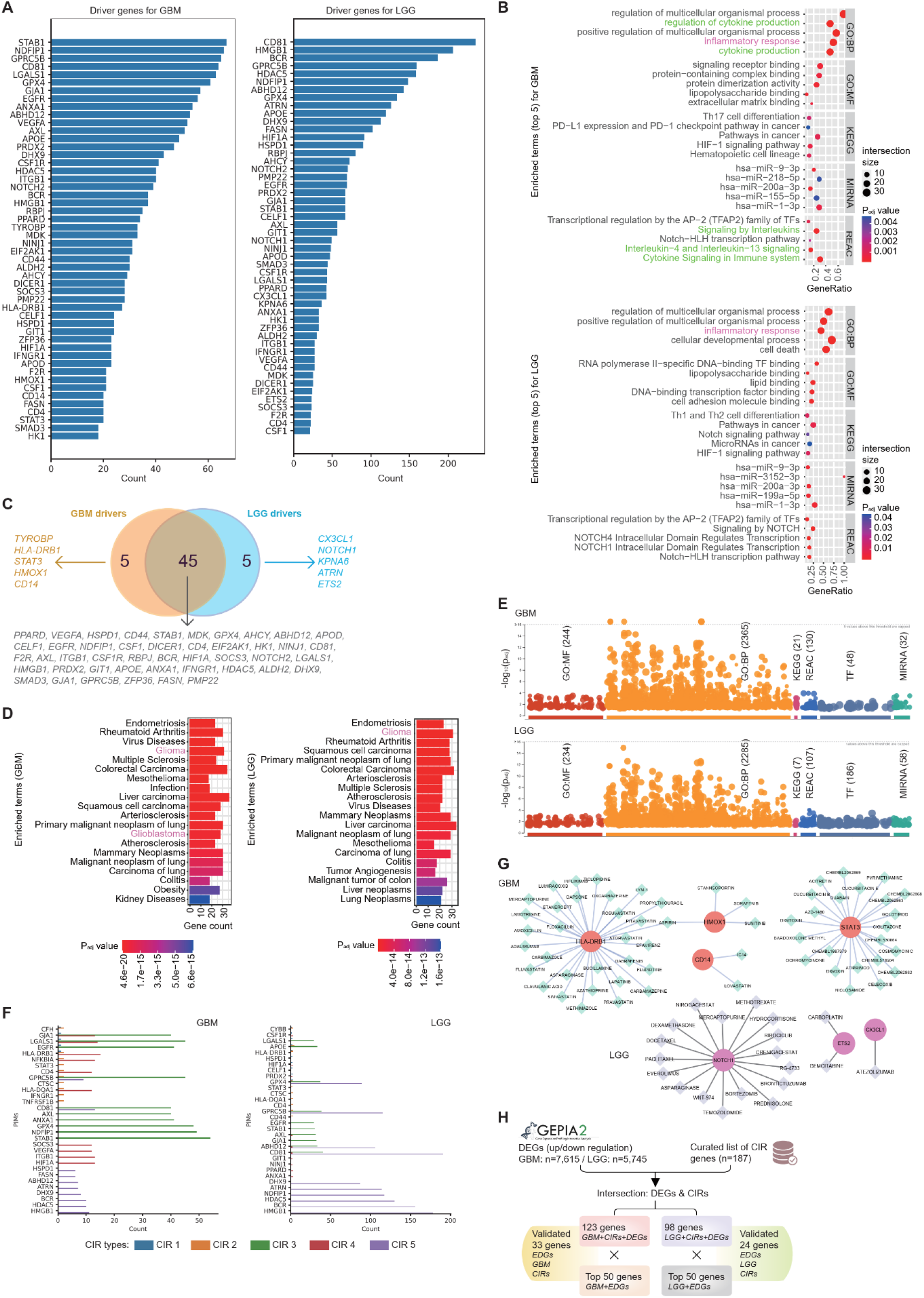
The identification of the essential driver genes (EDGs) to explain the TIME for glioma and its validation at population level. (A) The top-50 identified EDGs in GBM (left panel) and LGG (right panel) in desending order by frequency mapping. (B) Enrichment analysis of the top-50 EDGs in GBM (upper panel) and LGG (lower panel). (C) The intersection of the top-50 EDGs for GBM (left panel) and LGG (right panel). (D) Disease Ontology analysis of the top-50 EDGs in GBM (left panel) and LGG (right panel). (E) Gene set enrichment analysis statistics for the top-50 EDGs in GBM (upper panel) and LGG (lower panel) using Benjamini-Hochberg p-value adjustment method. (F) The frequency count distribution of the top-10 EDGs of each CIR type for GBM (left panel) and LGG (right panel). (G) The knowledge-based DGI network for the distinctly identified EDGs related to GBM (upper panel) and LGG (lower panel) revealing the significance of the discovery. (H) The stragegic validation model of the identified driver genes.

**Figure 4:**
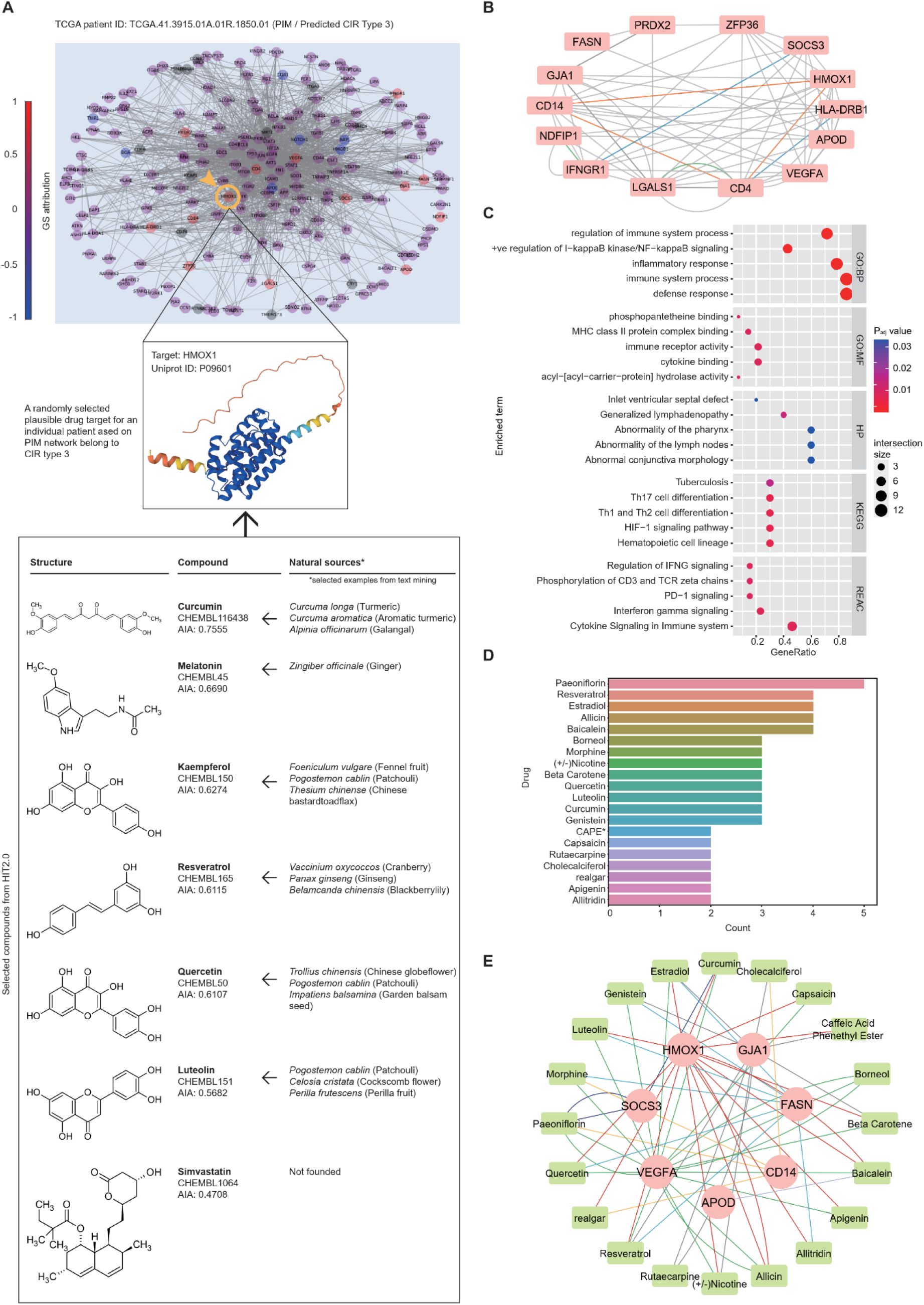
The personalized inflammatory mediator (PIM) network of an individual diagnosied with GBM and the proof-of-concept pipeline of personalized co-drug discovery. (A) The PIM network (upper panel) of a randomly selected GBM patient, and randomly selected one of the personalized effective driver genes (pEDGs), HMOX1, with its protein-level secondary structure which represens a plausible drug target to modulate the specific pEDGs, HMOX1 (mid panel); and the HMOX1-tagerting leads identified from the natural sources ordered as per their anti-inflammatory activity (lower panel). (B) The gene-gene networks for the 14 positive pEDGs using GeneMANIA. (C) The enrichment analysis of those 14 pEDGs. (D) The frequency count distribution of the top-20 leads ordered as per their anti-inflammatory activity. (E) The DGI network of the top-20 leads and their corresponding target genes indicating the personalized multiplexed drug-gene networks.

### 2.1. The XAI model characterizes five CIR types and outperforms the conventional NLR-based glioma prognosis

The proposed XAI model deciphered the PIM networks from the gene expressions (bulk RNA-seq of brain tissues) of the glioma patients from the The Cancer Genome Atlas (TCGA that consists of 165 GBM patients and 529 LGG patients (Figure 2A). The PIM networks characterize the CIRs associated with glioma patients into five distinct types. We obtained a total of five rationalized CIR types through Davies–Bouldin index and Kruskal–Wallis one-way analysis of variance method at p-value<0.00001. They showed improvised prognostic assessments in terms of OS compared to the conventional NLR-based prognosis at p-value<0.0001.

The PIM networks constituted inclusive CIR-associated molecular genetic information from the curated resources such as GO, DisGeNet, Biocrata, and Pubmed (supplementary S1). After pre-processing the data, we used 187 genes to enrich the Gene Set Variation Analysis (GSVA) and found nine significantly enriched CIRs-related pathways (Figure 2B, left panel; see supplementary 3.1). We identified the subsequent regulatory transcriptional factors (TFs) for each of those nine pathways for GBM and LGG (Figure 2B, right panel). Each column of the heatmap represented each patient, and the color map indicated their enrichment score across those nine pathways. The typified CIRs were stated in the upper panel of Figure 2B. Then, we have opted for the CIR type 3 and type 5 for the GBM and LGG, respectively in the successive analyses because of their significant abundance compared to the other CIR types (Figure 2F).

We showed that the CIR type 2 and type 4 were highly inflamed and had poor OS for GBM, and LGG, respectively. OS was found to be significantly lower in those with a high level of CIRs, and vice versa. The typified CIRs offered improvised prognostic efficiency for glioma patient subpopulations compared to the conventional NLR-based methods (Figure 2D-E). Besides, the NLR-based prognosis approach constantly faces multiple challenges(11) including the limitations beyond informing the all-or-none status of the CIRs (Figure 2E). It could only inform whether any inflammatory activity occurs in the local tumor environment which is also not statistically significant (Figure 2E). On the contrary, we showed that the data-driven typification of CIRs could be a better option as it explains higher-resolution insights about the patient-specific CIRs, related molecular pathways, and transcriptional controls (Figure 2B, 2D, 2G) with statistical significance. So, the demonstrated XAI model to typify the CIRs was data-driven, rational, and explainable that also outperforms the conventional prognostic assessment for glioma.

### 2.2. The XAI model identifies EDGs and explains the glioma TIME

We identified the EDGs that critically determine the TIME for the GBM and LGG patients irrespective of the CIR types. The top-50 EDGs (descending orders) for GBM and LGG were shown in Figure 3A. We found 45 genes in the intersection, and for GBM and LGG, we found exclusively five genes for each (Figure 3C). In addition, the CIR-group-wise abundance of identified EDGs (top-10 for each group) was shown in Figure 3F. We analyzed the drug-gene interactions (DGI) for those GBM- and LGG-specific EDGs (Figure 3G, and supplementary 3.2). Interestingly, the enrichment analyses using the glioma-wide EDGs showed that they were significantly involved in inflammatory responses (Figure 3B, highlighted in pink) for both GBM and LGG. Moreover, the GBM-specific EDGs were also found to be involved in inflammatory response-related biological processes such as cytokine production and its regulation (Figure 3B, highlighted in green), and also some inflammatory response-related pathways such as interleukin and cytokine signaling (Figure 3B, highlighted in green). Conversely, LGG-specific EDGs were significantly involved in an inflammatory response and cancer-related pathways such as NOTCH signaling (Figure 3B, lower panel). All the selected enrichment terms were statistically significant at p-value<0.005 for GBM (Figure 3B, upper panel) and p-value<0.05 for LGG (Figure 3B, lower panel). The entire enrichment terms are graphically shown in Figure 3E, and descriptions are in the supplementary S2. Next, the disease ontology (DO) analysis using DisGeNet in Enrichr showed that those EDGs were enriched in glioma (Figure 3D) at a p-value<0.0001. Interestingly, the GBM-specific EDGs also showed a distinct DO enrichment in glioblastoma (Figure 3D, left panel) at a p-value<0.0001.

Hence, the enrichment analyses validated the identified EDGs discovered by our XAI model to be relevant in the inflammatory response and glioma context. It also revealed the glioma TIME-specific molecular pathways and plausible regulatory molecules such as TFs and miRNAs, and the explored empirical associations further strengthen our analyses. Altogether, the analyses in Figure 3 supported the rationale behind the construction of PIMs and their deployments, including rationalizing the CIR types at the glioma-wide population, suggesting that the PIMs are explainable in terms of population-wise GO and DO enrichment.

### 2.3. Validation of the EDGs for glioma identified at population-level

The constructed pool of benchmarked differentially expressed genes (DEGs) related to CIRs and glioma validated the identified EDGs at population level (Figure 3H). A total of 123 and 98 genes were found to be differentially expressed in GBM and LGG, respectively, which are also linked to CIRs. Furthermore, those DEGs were validated against the top-50 EDGs for GBM and LGG. Interestingly, 66% of the EDGs (33 out of 50) were validated to be DEGs related to CIRs in GBM, wherein 48% of the EDGs (24 out of 50) were validated for LGG (Figure 3H). The validated GBM-specific EDGs were found to be enriched in inflammatory responses, cytokine signaling, and interleukin (IL-4, IL-13) signaling pathways and also showed enriched phenotypes such as lymphadenopathy, abnormality of the lymph nodes, and memory impairments (supplementary S2). The validated LGG-specific EDGs were enriched in inflammatory responses, cytokine signaling, NOTCH signaling, and interleukin signalling pathways. So, the identified EDGs and the validated EDGs showed consistent functional enrichments (Figure 3H, and supplementary S2).

### 2.4. The XAI model strategizes the personalized co-drug discovery to modulate the glioma-linked CIRs

The XAI-driven construction of PIM networks is also explainable at the individual level alongside the rationalization of the CIR types at the glioma-wide population. It helps in identifying the potential targets for modulating the patient-specific CIRs and support co-drug intervention. We constructed a total of 694 PIM networks from the total patient population (Supplementary S1). A group of EDGs driving the decision for PIM network construction were identified (Figure 4A). The GradientShap (GS) algorithm characterized those EDGs into four classes. One, the driver genes (red-marked); two, the non-driver genes (purple-marked); three, the ambiguous genes (blue-marked); and four, the knowledge-based gene lists aligned with the STRING database (grey-marked) within each PIM network. The set of ambiguous genes does not contribute to deciding the network toward its functional output, such as inflammatory response; instead, they negatively interfere in the decision-making process for the driver genes. So, in our analyses, we considered those pEDGs as they could be further validated for DGI target identification to modulate the CIRs. Next, we investigated the characteristics of those pEDGs in each PIM network, and the gene-gene interactions (GGI) were mapped (Figure 4B, and supplementary S3.3).

Next, we demonstrated our results with a single GBM patient (TCGA Patient ID: TCGA.41.3915.01A.01R.1850.01) belonging to the identified CIR type 3. The patient was not under any chemotherapy or radiotherapy at the point of sample collection as per the TCGA clinical history. The XAI model identified 14 pEDGs in that patient (Figure 4A, upper panel). The functional enrichment analyses of those 14 pEDGs showed significant involvement in inflammatory responses, defense responses, immune system processes, and regulations (Figure 4C). They were substantially linked to several inflammatory and inflammatory-linked cancer pathways, such as IFNγ signaling, regulation of IFNγ, cytokine signaling, PD-1 signaling, and different IL-signaling (IL-4, IL13). Besides, the Human Phenotype (HP) enrichment showed their linkage to different phenotypes, such as lymphadenopathy and abnormality of the lymph nodes, which indicates their involvement in metastasizing the primary tumor GBM through lymphatic systems(16). Also, abnormal inflammatory responses, vasculitis, pericarditis, respiratory tract infection, pulmonary infiltrations, abnormality in B-cell physiology, and humoral immunity with impaired increased inflammatory responses were highly linked phenotypically (Supplementary S2).

Next, to demonstrate the co-drug discovery to manage the concurrent inflammatory responses at the individual patient level, we showed the strategy with the above patient as example. Of those 14 pEDGs, we randomly selected a gene, *HMOX1*, and showed the associated co-drug discovery strategy. We used Herbal Ingredient’s Target version 2.0 (HIT2.0)(17) database to identify the related herbal drug targets against those 14 pEDGs. The active compounds and associated natural sources were identified (Figure 4A and 4E), and ordered as per their anti-inflammatory activity (AIA) scores(18). Our results suggested that the active compounds, such as curcumin, melatonin, kaempferol, resveratrol, quercitin, luteolin, and simvastatin, could potentially target HMOX1 to partially modulate the CIRs to the concerned patient. Curcumin possessed the highest AIA among them, whereas simvastatin had the lowest (Figure 4A). The natural sources for the active compounds were shown (Figure 4A, lower panel). For example, turmeric and galangal are the prominent resources for curcumin.

The GGI network was constructed among those 14 pEDGs (Figure 4B, and supplementary S3.3). They were enriched in the following biological processes: regulating the immune system processes, inflammatory responses, and NFkB signaling; pathways: IFNγ signaling, PD-1 signaling, and HIF-1 signaling; and phenotypes: lymphadenopathy, abnormal lymph nodes, and ventricular septal defects (Figure 4C). Then, the anti-inflammatory natural compounds targeting those 14 pEDGs to modulate the CIRs of the individual, we constructed an ordered frequency map with the interacting compounds against their corresponding target pEDGs. The top-20 ordered compounds were listed in Figure 4D. We also mapped the DGIs of those top 20 compounds and their target pEDGs (Figure 4E).

So, the XAI model analyzed the single-patient transcriptomic data to indicate a critical notion in personalized medicine. Individual PIM network analyses facilitate strategizing the co-drug discovery from natural sources to modulate individual CIRs. So, the XAI model supports the proposed co-drug discovery capacity.

## 3. Discussions

Managing CIRs is one of the strategic areas in the next-generation synergistic co-drug discovery in oncology. Thise strategic focus can be beneficial as they are engineered to be non-interfering with the primary chemotherapy but will function synergistically to modulate the concurrent CIRs.

Our proposed XAI model uniquely characterizes the molecular-genetic, personalized, and disease-level facets of CIRs to typify them. This strategy leveraged to recommend suitable natural compounds-based interventions to alleviate personalized CIRs. However, such validation is one of the key bottlenecks in personalized medicine. Interestingly, our model provides an individual-level CIRs information that could flexibly be scaled up to the population level, offering a verifiable scenario of PIMs at the disease level. Further the typified CIRs indicated an improvised prognostic assessemt for glioma and supported the co-drug discovery to combat the unnecessary CIRs.

### 3.1. The XAI-based typified CIRs offer an enhanced molecular characterization of glioma patients that outperformed the conventional NLR-based prognosis

The typified CIRs indicated an enhanced prognostic assessment for glioma through rationalizing its TIME. The trained XAI model facilitated understanding the CIRs from the deep molecular and cellular perspective to synthesize an explainable and rationalized insights. It served as the basis for our CIR typification. Our model identified the PIMs and converged the information at the disease level, and allowed disease-level phenomena discovery to deploy back in a personalized manner. GBM, being the most lethal type of glioma shows nearly 3-5% OS rate. Its anatomical occurrence and complex TIME made its diagnosis and prognosis challenging(19). Frequently occuring vascular edema, increased intracranial pressure and leakage in blood-brain barrier impedes the prognostic and therapeutic management in glioma, especially in GBM patients(20–22). It turns GBM more aggressive and resistant to the treatments. Interestingly, we reported that the inflammatory responses, lymphadenopathy, abnormal lymph nodes and vasculature were the most enriched biological processes and phenotypes discovered in the GBM population (Supplementary S2, and Figure 3H). Impaired cerebral lymphatic systems were reported to trigger such cerbral vascular edema(23). A study on a rat stroke model showed that brain injury might activate peripheral immune cells and aggravate CIRs plausibly by activating tyrosine kinase pathways(24). The VEGF-C/VEGFR3 signaling was identified as the underlying pathway for activating those responses. And inhibition of VEGFR3 reduces lymphatic endothelial activation and minimizes it(24). Thus, we may consider that the unresolved CIRs could cause abnormality in the cerebral lymphatic systems that might produce cerebral vascular edema. And the under-characterized influences of the CIRs may turn the glioma, especially the GBM, more resistant to the treatment with a poor prognosis.

Temozolomide (TMZ) is widely used as standard-of-care to treat newly diagnosed glioma(4,25), and sometimes in combination with adjuvant immunotherapy(26). VEGF inhibitors (27), and later, bevacizumab (BVZ), a monoclonal antibody was approved for GBM treatment(28). However, the recurrence of GBM remained a major constraint(29–31). Besides, TMZ exerts immunomodulatory effects on the TIME. They have been reported to be involved in two contradicting circuits, one, disrupting the local immunosuppressive mechanisms, and two, exerting detrimental influences on the peripheral immune response(26). Later one drives the treatment failure(26). Next, the programmed death-ligand 1 (PD-L1) critically regulates the immune responses and often prevents exacerbated activation and autoimmunity. So, the PD-L1 overexpressing tumors often show poor prognosis, and the TMZ was reported to upregulate PD-L1 contributing to the poor prognosis of glioma and may influence the treatment failure due to induced immunosuppressive activity(32). Interestingly, in our results, the typified CIRs in GBM population identified PD-L1 expressions and PD-1 checkpoint pathways as one of the most enriched pathways. As per the TCGA clinical data, mostly the GBM patients received TMZ as primary chemotherapy. So, we may perceive that the aggravated CIRs could stimulate PD-L1 overexpression in GBM patients making them more resistant to the treatment with a poor prognosis (Figure 3B). It certainly prompts to investigate the individual PD-L1 profile to strategize the therapeutic planning for GBM patients.

Our results offered a unique advantage over NLR-based prognosis in OS assessment (Figure 2D, 2E). The typified CIRs allowed us to pay precise attention to the subsets of patients and even at the personalized level that can help to customize the prognostic assessments. To achieve this, identifying the CIRs-related EDGs is one of the key demands in oncology which we addressed here.

### 3.2. PIMs explain the TIME for glioma at population-level

We identified the glioma TIME-determining EDGs. The enrichment analyses of those EDGs at the disease level were significant to endorse the relationship between the inflammatory responses, related pathways, and their associations. The DGI for those GBM- and LGG-specific EDGs were also consistent with the past studies (please see the result section 2.2 and supplemenatry S3.2). For example, a GBM-specific distinct driver gene, *HMOX1*, was involved and elevated predominantly (>90%) among the GBM patients, confirmed by immunohistochemistry analysis(32). Interestingly, they also reported the involvement of *HMOX1* in stemness of GBM and studied it as a putative cancer stem cell markers(32). This is exciting as it is consensus with our identified GBM-specific EDGs, *HMOX1*. Another study targeted *HMOX1* for engineering an exosomal delivery system loaded with STAT3-specific siRNA for synergistic therapy against TMZ-resistant GBM(33). This *in vivo* synergistic therapy decreased tumor proliferation by inhibiting the expression of *MGMT*. So, this finding strongly advocates our rationale to identify the data-driven, evidence-based targets for co-drug discovery for modulating the hidden CIRs. Excitingly, the *HMOX1*, suggested by our XAI model was strongly rationalized for the synergistic therapeutic targets in GBM (Figure 3C). Another study showed that *STAT3* was designated to stratify glioma patients for targeted therapy(34). A higher abundance of *STAT3* was reported in the aggressive form of glioma, and the GBM, and *STAT3*-low tumors were reported in LGG. We also identified *STAT3* as another key GBM-driving EDGs (Figure 3C). Next, *CX3CL1* was reported to be involved in the progression of LGG(35) which is consistent to our results (Figure 3C). We also reported *NOTCH1* as another key LGG-specific EDGs (Figure 3C), but the related NOTCH-signaling pathway still remained deceptive in distinguishing the glioma grades and underlying inflammatory response connections(36). It certainly stimulates a scope of investigation to justify the role of NOTCH1 in LGG and GBM and to devise it as a therapeutic target.

The identified IL-6/JAK/STAT3 pathway was significantly activated in the GBM patients, especially with CIR type 3 and type 4, the two most highly inflamed GBM subgroups (Figure 2B). The IL-6/JAK/STAT3 pathway was identified in different cancers. Increased IL-6 was reported to be associated with high CIRs and complex diseases such as rheumatoid arthritis, inflammatory bowel disease, and solid tumors, including GBM(22,37). Our results specified the activation of IL-6/JAK/STAT3 in the CIR type 3 and 4 in GBM but not in the most dominant CIR type 5 of LGG, which strengthens the claim that those two highly inflamed groups of GBM patients showed higher activation of IL-6/JAK/STAT3 pathway resulting poor prognosis (Figure 2C, upper panel). Interestingly, we also observed that the second largest subset of inflamed LGG, the CIR type 3, showed apparently higher activation of IL-6/JAK/STAT3 and resulted in poor prognosis among the LGG cohorts (Figure 2C, lower panel, blue-colored area). So, the prognosis among the GBM and LGG patients could be determined by that specific pathway and vary across the typified CIR groups.

Altogether, our results showed that the identified EDGs explained the glioma-specific TIME. The XAI-based identified EDGs outperformed the DEG-based hub gene identification method as it is limited to deploy in a personalized context. DEGs are inclined at the population level. So, they may serve as the disease-level biomarker where the minimum effective population should be more than 10 to be statistically significant. On the contrary, the XAI model identified the EDGs from personalized data which can also be scaled at population.

### 3.3. PIMs explain the individualized glioma-specific TIME and support personalized co-drug discovery to modulate the CIRs

The XAI-derived PIM networks explained the glioma-specific TIME at individual patient alongside the rationalization of the CIR types at the population. It helped to identify the prospective targets for optimizing the patient-specific CIRs through synergistic co-drug discovery. Conventionally, the DEGs serve as the benchmark for identifying group-wise gene features, demonstrating their effectiveness in many studies. And in terms of individuals, the existing methods for identifying DEGs, for example, DESeq2, limma, etc., are only effective in a sample size >10(38). So, we need a method that could be effective at an individual level. Here, we deployed the GS algorithm to identify the pEDGs representing individual-level gene features (Figure 4A). The selected example was not under any chemotherapy, so the probability of chemotherapy-induced CIRs can be ruled out.

Further, we exemplified one of the pEDGs, *HMOX1*, which has been prioritised in many GBM therapies, especially TMZ-resistant GBM. This personalized analysis suggested a relevance to develop anti-inflammatory measurements to tackle abnormal lymphatic growth and vasculature in consistence with the enrichment data (Figure 4C) and prior discussions (Section 3.1, and 3.2) to minimize the chances of cerebral vascular edema, blood-brain barrier leakage and induced brain injury in glioma. In this view, co-drug discovery is essential to synergistically augment the primary chemotherapy’s responsiveness while managing the concurrent CIRs. Significance of several XAI-identified pEDGs as potential targets for optimizing CIRs in GBM has been exphasized in several studies(39,40). The top-ranked co-drug, Paeoniflorin, was shown to interact with *SOCS3* in the specific patient’s PIM. Paeoniflorin was reported to upregulate *SOCS3*, a well-known inflammation brake, by inhibiting the ASK1-TF axis and reducing inflammation(39). Several other studies suggested Paeoniflorin’s impact in inhibiting EMT and angiogenesis in GBM in humans via K63-linked c-Met polyubiquitination-dependent autophagic degradation(40), and by suppressing TGFβ signaling pathway(41). Interestingly, the EMT pathway was shown to be highly activated among the CIR type 3 for the GBM patients (Figure 2B), wherein the type 3 CIR was indicated as the most dominant GBM-related CIR type (Figure 2F). The CIR-induced EMT activation may contribute to poor prognosis in GBM. Hence, hypothetically, we may recommend that patient to take Paeoniflorin as a co-drug along with their primary chemotherapy. The top-ranked compounds did not show any contraindication to the TMZ using WebMD, RxList, and DrugBank.

### 3.4. The XAI model rationalizes the personalized findings at the population level and vice versa

Our XAI model enabled switching between the population-based and personalized parameters from the omics data. Personalized medicine is often limited by a largely unexplored application of high-throughput omics data from the population study. However, our XAI model showed an efficient way of transitioning from population to individual level and vice versa. It may have significant consequences for improving patient outcomes and prognosis.

The population-level bulk omics analyses often mask the heterogeneity within the EDGs expression across individuals. So, offering accurate information for personalized medicine, it is necessary to have a meticulous analysis of individual data. Here, our XAI model enables analyzing the single-patient transcriptomic data, which shows a critical path in personalized and precision medicine. PIM network analyses facilitate strategizing the co-drug discovery from natural sources to modulate individual CIRs. So, the XAI model extends the drug discovery capacity to switch between population-based, subset-based and personalized contexts.

## 4. Conclusions

We proposed a strategic manipulation of CIRs by reversely engineering the attributes of inflammatory mediators driving those respnses in glioma patients at personalized level using XAI model. It possibly widen an avenue to recognize the manisfestations of CIRs underlying the glioma development, progression, and treatment responsiveness. Our findings highlight a novel strategy to typify CIRs as prognostic markers for glioma patients. We also showed a systematic way to discover natural compund-based co-drugs for ensuring better survival upon optimizing the inflammatory responses developed as a counterpart in the course of glioma progression and therapy.

## 5. Methods

We proposed an XAI model to redefine CIRs as prognostic markers for glioma using 694 patients’ data derived from TCGA. First, we identified CIR-related genes from the five curated databases, and nine HALLMARK pathways were enriched using gene set variation analysis (GSVA). We clustered them as per signature CIRs. Then, we trained the XAI model to systematically characterize the glioma-specific PIMs using multifaceted regulators such as TF networks, and cellular infiltration markers. It returned rationalized typified CIRs whose relevance as PMs was validated using OS analyses. Then we showed a personalized co-drug strategy for modulating CIRs. The detailed study scopes, criteria, and considerations were discussed in the supplementary S3.4.1 and 3.4.2.

### 5.1 Data description, analysis and pre-processing

The RNA-seq data from TCGA-based transcriptome were obtained using TCGAbiolinks with R (http://bioconductor.org/packages/TCGAbiolinks/)(42) as STAR raw counts. The primary and recurrent solid tumor samples from 165 GBM patients and 529 LGG patients were analyzed in this study. In pre-processing, we filtered out the possible outliers of patient samples using the *TCGAanalyze_Preprocessing* function. We defined a square symmetric matrix of Pearson correlation among all patients with cut-off value as 0.6. The samples with correlation cut-off <0.6 were outliers. Next, we filtered out the low-expressed genes using variance filtering function, *TCGAanalyze_Filtering* with default cut off as qnt.cut = 0.25 and var.cutoff = 0.75.

### 5.2. Clustering of the typified CIRs

GSVA enrichment output were clustered using a hierarchical clustering with ward distance. Patients with similar GSVA profile was considered within the same cluster (supplementary 3.4.3). The optimal clusters number was estimated using Davies-Bouldin evaluation and the *t-test* of the GSVA enrichment.

### 5.3. Construction and training of the XAI model

The XAI model construction consists of two parts; one, a convolutional neural network (CNN) model to train the dataset, and two, the GS algorithm to reversely engineer the significant gene set of each individual. We constructed the CNN model with six 2D convolution layers (CLs) with input channel set as 1 and the output channels set as 8, 16, 32, 64, 128, 256. Each CL followed a ReLU activation and at the second, fourth, and sixth layers, there were 2D max pooling layers(43) with the size of (1×3). Then, nine fully connected layers were applied. The out features of each fully connected layer were 1000, 400, 200, 128, 64, 32, 16, 8 and 5. Then, a softmax layer was used for generating the probability for each samples within different CIR types. The corresponding CIR type with the highest probablity was assigned to the sample. The initial learning rate was set at 0.0001 whereas the batch size and epochs were 256 and 30, respectively. And the adam optimizer was used for weight updating throughout the training process wherein cross-entropy loss was selected as the loss function.

After the training, we employed the GS algorithm(44) to decode a set of important genes for each patient. The algorithm details are mentioned in equation (1) and supplementary S3.4.4. It returned the final SHAP values represented as the expected values of gradients*(Inputs – Baselines). We used captum(45), a Python-based package to perform the GS algorithm.

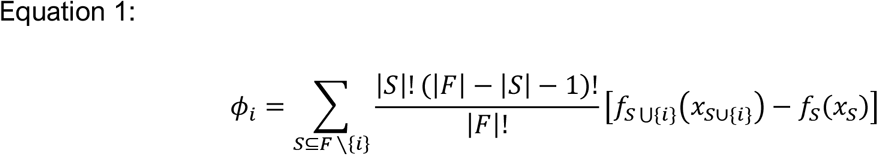

Where *F* represents full set of features, and *S* represents a subset of it. The *i* represents a specific feature in the set *F. f*_*SU*{*i*}_ is the model trained with feature *i*, and *f*_*S*_ is model trained without feature *i*.

### 5.4. Construction of the PIM networks

The XAI model deciphered a set of distinct inflammatory features for each patient. Relying on that, we constructed the PIM networks using STRING. The GS attribution helped to distinguish the CIR-driving EDGs (GS score > 0.1), non-EDGs (GS score -0.1 to 0.1), and the deceived proteins which deviates the decision making of the CIR-driving EDGs (GS score < -0.1).

### 5.5. Overall survival analyses

Prognostic significance of the XAI-based typified CIRs was demonstrated using OS analyses using Kaplan-Meier estimator. Log-rank tests were used for comparing overall survivor functions among the patients categorized within five CIR types. The OS-curves indicate the time to in-hospital death. The x-axis represents the elapsed time (in years) from the admission date, and the y-axis represents the survival probabilities.

### 5.6. Validation of the identifed EDGs at population and personalized level

To validate the GBM- and LGG-specific EDGs, we built a pool of benchmark DEGs linked to CIRs and glioma. It combined experimentally and computationally validated gene lists with a standard pan-cancer DEGs identifying tool, GEPIA2(46). The pool of DEGs was extracted using ANOVA with |Log FC >1| at p-value<0.01 from GEPIA2. Extraction of DEGs included both over- and underexpression of the genes, as the up/down-regulation was not a criterion for EDG identification. Then we quantified the pEDGs for each GBM and LGG patient using statistical frequency mapping. It helped to sort the top-50 (descending ranking) EDGs explainable in both contexts, personalized, and population. It also identified the higher frequency pEDGs.

### 5.7. Functional enrichment analyses

Gene ontology (GO)-enrichment analysis was used to determine the molecular functions (GO:MF) and biological processes (GO:BP). The Kyoto Encyclopedia of Genes and Genomes (KEGG), and Reactome Pathway Database (REAC) enrichment analysis were used to determine the cellular pathways of the identified genes. TFS and MIRNA enrichment analyses were used to determine the associated molecular regulators. Human phenotypes (HP) enrichment analysis was used to determine the enriched phenotypes associated with the identified genes. GO, KEGG, REAC, TF, MIRNA, HP were analyzed using g:Profiler(47). DisGeNet enrichment analysis was performed to determine the disease enrichment associated with the identified genes using Enrichr(48). The visualization of the enrichment analyses were executed using R.

### 5.8. Personalized co-drug discovery strategy

The personalized co-drug discovery pipeline was constructed with the identified PIM networks and the pEDGs. The pEDGs within each PIM network were considered as the potential target genes to modulate individual CIRs. We used the Herbal Ingredients’ Targets Platform (HIT 2.0) to search and curate the natural compunds-based leads against those targets. We retrieved the protein structures of those targets genes using AlphaFold(49). Next, we retrieved the chemical structures of natural compunds using ChEMBL and their corresponding anti-inflammatory activity (AIA) scores using InflamNat(18). Then, we ordered the compounds baased on the descending AIA scores. Later, we mapped the pEDGs and DGI networks using GeneMANIA(50) and Cytoscape (version 3.9.1) respectively. The g:Profiler was used to perform the functional enrichment analyses of those pEDGs. And we estimated the relevance and frequency of the identified candidate drugs for each PIM network using Python.

## Acknowledgments

Hong Kong General Research Fund (12102722, 12101018, 12102518, 12100719), Hong Kong Theme-based Scheme (T12-201/20-R), Interdisciplinary Research Matching Scheme Hong Kong Baptist University (RC-IRMS/15-16/01), Technology Innovation Strategy Special Fund (Guangdong-Hong Kong-Macau Joint Lab, No: 2020B1212030006) and Nature Science Foundation of Shandong province (ZR2020QH219).

