## Supplementary file for "An explainable artificial intelligence-based typification of chronic inflammatory responses enhances glioma prognosis"

**SUPPLEMENTARY FILES (S1 and S2):**

1. **Supplementary S1:** PIM networks for GBM and LGG (Accessible link: https://bit.ly/3kek6tG)
2. **Supplementay S2:** Validation output with GEPIA (Accessible link: https://bit.ly/3ICEbmP)

**SUPPLEMENTARY NOTE (S3):**

**S3.1. Results:**

The PIM networks constituted inclusive molecular genetic information associated with CIRs from different the curated resources such as GO (GO0006954, GO0002544), DisGeNet, Biocrata (Biocrata_Inflam_pathway), and Pubmed (PMID29929551). After pre-processing the data, we used 187 genes to enrich the Gene Set Variation Analysis (GSVA) and found nine significantly enriched CIRs-related pathways (Figure 2B). These are the apoptosis pathway, complement, allograft rejection, inflammatory response, interleukin (IL)-6-JAK/STAT3 signaling, hypoxia, epithelium-to-mesenchymal (EMT), tumor necrosis factor-alpha (TNFα) signaling via nuclear factor kappa B (NF-κB), and interferon-gamma (IFNγ) response.

We identified the subsequent regulatory transcriptional factors (TFs) for each of those nine pathways for GBM and LGG. For example, the apoptosis pathway was found to be transcriptionally controlled by three TFs such as HMGB2, RELA, and JUN (Figure 2B). The nine pathways were indicated in the left panel of Figure 2B, and the corresponding regulatory TFs were mentioned in the right panel. Each column of the heatmap represented each patient. The color map indicated their enrichment score across those nine pathways. In the upper panel, we stated the typified CIRs. We have selectively opted for CIR type 3 and type 5 for the GBM and LGG, respectively, because of their significant abundance compared to the other CIR types in the successive analyses (Figure 2F).

**S3.2. The XAI model identifies EDGs and explains the glioma TIME**

We analyzed the drug-gene interactions (DGI) for those GBM- and LGG-specific EDGs (Figure 3G). For example, the GBM-specific EDGs, *HMOX1*, shared sensitivity to the small molecule drugs sunitinib, an FDA-approved tyrosine kinase inhibitor[1,2], sorafenib, kinase inhibitor[3–6], and stannsoporfin, heme oxygenase (HO) inhibitor; and *CD14* was found to be sensitive to lovastatin, an HMG-CoA reductase inhibitor, and IC14, a CD14-specific monoclonal antibody. And the LGG-specific EDG, *NOTCH1*, shared sensitivity to the drug temozolomide (TMZ), the most commonly used chemotherapeutic drug for glioma[7] and other chemotherapeutic drugs such as docetaxel, paclitaxel, and methotrexate; and *ETS2* was found to be sensitive to chemotherapeutic drugs, carboplatin, and gemcitabine. Then, we investigated the empirical association among those identified EDGs and glioma types. For example, the relevance of GBM and the identified EDGs such as *HLA-DRB1*[8–11], *HMOX1*[12,13], *CD14*[14–16], *TYROBP*[17], and *STAT3*[18–20] were experimentally examined. On the other hand, the relevance of LGG and some of the identified EDGs such as *NOTCH1*[21,22], *ATRN*[22], *CX3CL1*[23–27] were experimentally studied. (all these RED marked part is to be placed in suppl.).

**S3.3. The XAI model strategizes the personalized co-drug discovery to modulate the glioma-linked CIRs**

The GGI network was constructed among those 14 pEDGs (Figure 4B). In that network, *CD4*, *CD14*, and *HMOX1* were found to be co-localized (Figure 4B, orange-colored edges); physical interactions were found between *CD4* and *LGALS1* (Figure 4B, green-colored edges); *SOCS3* and *IFNGR1*, and *CD4* and *HLA-DRB1* were found to be related at pathways level (Figure 4B, blue-colored edges) and rest other genes were co-expressed (Figure 4B, grey colored edges).

**S3.4. Methods:**

**S3.4.1. Consideration on glioma types:** Glioma includes both glioblastoma (GBM) and lower-grade glioma (LGG). In our study, all the lower grade (I-III) glial tumors such as astrocytomas, oligodendrogliomas, oligoastrocytomas, and ependymomas are referred to as lower-grade gliomas (LGG), and the grade IV as glioblastomas (GBM)[28]. The data pre-processing step to filter out the possible outliers of patient samples using the *TCGAanalyze_Preprocessing* function performed an Array Array Intensity correlation (AAIC).

**S3.4.2. Criteria, considerations, and study scopes:** This study focused on typifying the CIRs associated with glioma. We encompassed inclusive CIRs linked patients diagnosed with glioma. It included the CIRs across all phases involving glioma development, progression, and during the chemotherapy to the patients. So, we set our inclusion criteria for the subjects that strictly relied on the disease label (as GBM and LGG) at the point of the primary diagnosis of those patients. And we intended to investigate the wide-ranging CIRs from the perspective of inflammatory mediators linked to all those phases. As we prioritized improvising the glioma prognosis through optimizing the CIRs. Thus, the scopes of our study do not rely on the glioma patient-specific clinical and demographical variables. So, our constructed XAI model can be applied irrespective of the time of intervention from the point of diagnosis and the type of intervention given to the patients.

**S3.4.3. Anchoring the TCGA patient data to the CIR-related curated functional signatures:**

To establish the first-line relationship among the TCGA patient data and the CIRs, we employed a data mining strategy with the curated molecular genetic signaure endorsed for sharing CIR features. Then, the single-sample Gene Set Enrichment Analysis (ssGSEA) was performed to estimate the enrichment score of each patient using the R package 'GSVA'[29] to identify up- and downregulated genesets of interests or pathways within GBM and LGG. The CIR-related molecular genetic signature was obtained from curated sources such as GO (GO0006954, GO0002544), DisGeNet, Biocrata (Biocrata_Inflam_pathway), and Pubmed #PMID29929551. And the hallmark genesets were obtained from the Molecular Signatures Database (MSigDB, V7.2)[30].

**S3.4.4. Construction and training of the XAI model:**

After the training, we employed the GS algorithm[31] to decode a set of important genes for each patient. The algorithm details are mentioned in equation (1). It estimated SHAP values by computing the expectations of gradients through random sampling from the distribution of baselines/references. It introduced a white noise to each input sample (n_samples*n_times), and selected a random baseline from the distribution of baselines and a random point along the path between the baseline and the input, to compute the gradient of outputs as per those selected random points.

We constructed the PIM networks using STRING[32].
